# DNA barcode-guided lentiviral CRISPRa tool to trace and isolate individual clonal lineages in heterogeneous cancer cell populations

**DOI:** 10.1101/622506

**Authors:** Y Akimov, D Bulanova, M Abyzova, K Wennerberg, T Aittokallio

## Abstract

The genetic and functional heterogeneity of tumors imposes the challenge of understanding how a cancer progresses, evolves and adapts to treatment at the subclonal level. Therefore, there is a critical need for methods that enable profiling of individual cancer cell lineages. Here, we report a novel system that couples an established DNA barcoding technique for lineage tracing with a controlled DNA <u>b</u>arcode-<u>g</u>uided <u>l</u>ineage <u>i</u>solation (B-GLI). B-GLI allows both high-complexity of lineage tracing and effective isolation of individual clones by CRISPRa-mediated induction of puromycin resistance, making it possible to unbiasedly trace, isolate, and study individual cancer cell lineages. We present experimental evaluation of the system performance in isolation of lineages and outline a comprehensive workflow for B-GLI applications. We believe the system has broad applications aimed at molecular and phenotypic profiling of individual lineages in heterogeneous cell populations.

## Introduction

Cancer consists of multiple genetically related, distinct subclones ^1^ that evolve in parallel and display heterogeneity at genomic^2,3^, epigenetic^4–7^, or phenotypic level^8–10^. As malignancy progresses, particular subclonal populations of cancer cells develop differing aggressive phenotypes, such as faster proliferation, metastatic potential^11,12^, or drug resistance^13,14^. To understand which molecular features promote an aggressive phenotype in particular subclones, and hence, further the tumor progression, elucidating the biology of individual cell lineages would be particularly insightful.

DNA barcoding (tagging of individual cells with unique DNA sequences) enables high-throughput tracking of individual clonal lineages by next-generation sequencing (NGS). The barcoding approach accurately and quantitatively determines the lineages that get enriched or depleted following tumor initiation^15,16^, drug resistance^13,17–19^, metastatic spread^11,12,20^, or another type of phenotype selection. Moreover, current developments allow for the use of DNA barcoding to link multiple phenotypes to a lineage^21^, or even to couple lineage tracing with single-cell transcriptomics^22^. A number of DNA barcoding-based methods have been developed to date^13,23–27^, but the limitation of NGS-based applications is that they are cell-destructive *per se* and prohibit isolation of subclones for further functional or molecular profiling. Therefore, there is an unmet need for robust methodologies that couple high-throughput lineage tracing with controlled isolation of the individual lineages.

The current approaches to study individual subclonal lineages have several shortcomings inherent to the methodology. Lineage tracing using classical fluorescent tagging allows one to isolate tumor-initiating subpopulations^28^, but it has a low resolution for studying complex populations due to the very limited number of fluorophores that can be studied simultaneously. A manual subcloning of single cancer cell cultures followed by molecular and phenotypic profiling of the clones has yielded important insights into the biology of metastatic^12^ and chemoresistant subclonal lineages^19^. However, the throughput of manual cloning is medium-to-low and the approach requires prolonged cell culture that may change cloned cells over time^29^. Similar limitations apply to functional selection, such as selection of drug-resistant subpopulations by drug exposure^30^ or selection of metastatic cells by *in vivo* invasion assay^31^. These approaches allow for limited understanding of whether the drug-resistant or metastatic clones pre-existed in the population or developed the phenotype *de novo*, and if so, what were the determinants of phenotypic plasticity underlying the clonal selection.

To address these challenges, we developed B-GLI - a novel lentiviral high-throughput lineage tracing system that enables one to isolate lineages of interest from the bulk cell population by their DNA barcodes. Unlike the existing DNA barcoding libraries, we designed the barcodes to serve as recruitment sites for the dCas9-fused transcriptional activator to guide CRISPR activation of the puromycin resistance in the specified lineages in a highly-controlled manner. We demonstrate the specificity of the method in an validation experiment that modelled the isolation of prevalent subclonal lineages (∼0.1% of population) that displayed a distinct rare phenotype in pancreatic cancer cells. We also provide a comprehensive protocol and vectors for public use of B-GLI in applications aimed at phenotypic characterization of cancer subclones.

## Results

### Barcode-guided lineage isolation (B-GLI) strategy

To combine DNA barcoding with the isolation of specific lineages, we devised a lentiviral system that functionalize DNA barcodes to serve both as heritable cellular tags for monitoring clonal dynamics and as recruitment sites for the catalytically inactive Cas9 (dCas9)-fused transcriptional activators. For that, we placed DNA barcodes containing protospacer adjacent motifs (PAMs) upstream of the basally-inactive promoter controlling expression of puromycin resistance gene. Lineage-specific activation of the selection marker upon targeting the barcode with sgRNA provides accurate means for selecting a lineage based on its identifier - the sequence of the DNA barcode (Figure 1A).

**Figure 1.**
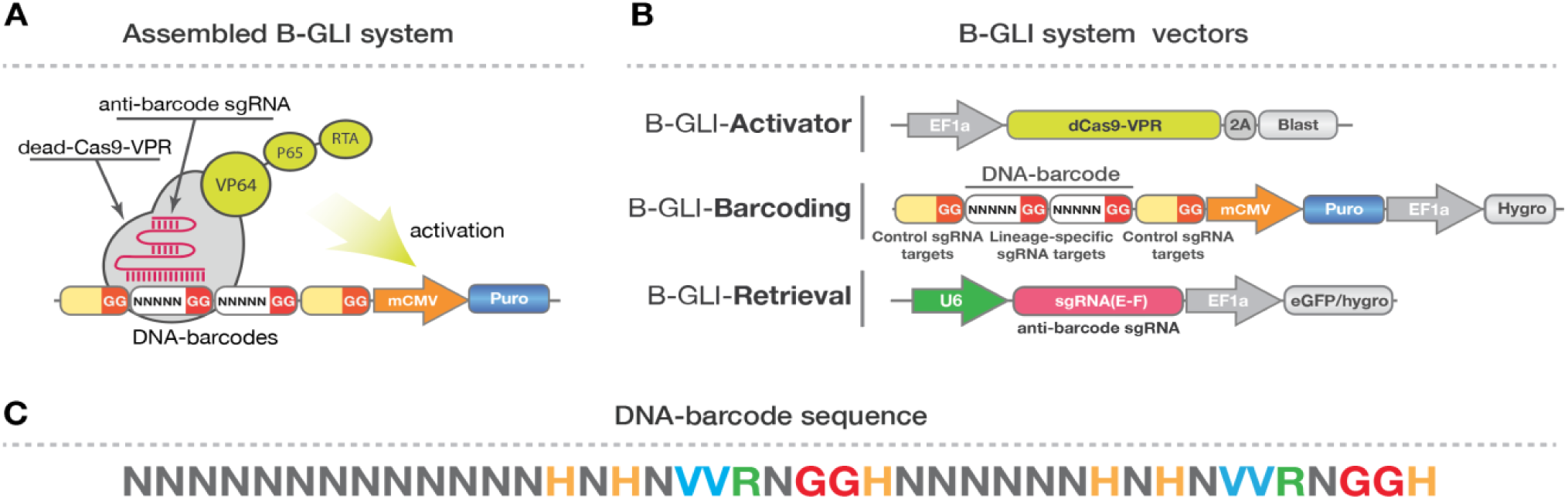
The functional components of barcode-guided lineage isolation (B-GLI) system. (**A**) Assembled B-GLI system. A sgRNA against a specific DNA-barcode forms a complex with a dCas9-VPR protein. A complex binds to the DNA-barcode region upstream of mCMV promoter only in a lineage with a barcode complementary to the sgRNA. Binding of the sgRNA-dCas9-VPR complex activates the expression of puromycin from the otherwise inactive mCMV promoter. (**B**) Graphical representation of the B-GLI lentiviral vectors. EF1a, elongation factor 1α promoter; dCas9-VPR, catalytically inactive Cas9 protein fused to VP64-P65-RTA transcriptional activator; 2A, 2A self-cleaving peptide; Blast, Blasticidin resistance marker; mCMV, minimal CMV promoter, Puro, Puromycin resistance marker; Hygro, hygromycin resistance marker; U6, human U6 RNA polymerase III promoter; sgRNA(E-F), single guide RNA cloning site and optimized scaffold. eGFP, enhanced green fluorescent protein. To simplify the illustration, the following common lentiviral elements are not shown: Rev response element, central polypurine tract, Psi packaging signal, Flag octapeptide tag, Woodchuck Hepatitis Virus Posttranscriptional Regulatory Element. (**C**) Design of DNA-barcoding sequence, where degenerate bases were chosen to maximize the efficacy of sgRNA targeting. Single letter abbreviations are as follows: N = A, C, G or T; H = A, C or T; V = A, C or G; R = A or G.

To implement the barcode-guided isolation strategy, we constructed 3 lentiviral vectors (Figure 1B; please see Methods and Supplementary Files for the details of the cloning experiments). The B-GLI-Activator vector carries dCas9 fused to a highly-effective transcriptional activator domain VP64-p65-RTA (VPR)^32^. The B-GLI-Barcoding vector carries a semi-random 37 bp DNA barcode that is interspaced by 2 PAMs, a basally-inactive minimal CMV(mCMV) promoter, and a puromycin resistance cassette. The barcode region was designed to maximize the efficiency of dCas9 binding according to the previously described nucleotide preferences for sgRNA efficacy^33^. The barcode region also includes “mock control barcodes”, sequences with common sgRNA target sites, which allows for activating puromycin resistance marker expression in all the barcoded cells (mock control). Finally, B-GLI-Retrieval vector delivers the barcode-targeting “anti-barcode” sgRNA that directs dCas9-VPR activator to the mCMV promoter and puromycin resistance cassette. Therefore, when the barcoded cells receive the B-GLI-Retrieval vector, the puromycin resistance gene expression will be activated only in the lineages with matching sgRNA-barcode pair (Figure 1). To further improve the efficiency of transcriptional activation, we implemented the previously optimized sgRNA(E-F) scaffold design, which has been reported to improve dCas9-sgRNA complex binding to the DNA^34^.

### Experimental workflow using B-GLI

An experiment aiming to identify and isolate phenotypically distinct lineages (e.g. those displaying drug response, differentiation to a particular type of cells, or invasion to a particular organ in vivo) using B-GLI system includes the following steps (Figure 2):

**Figure 2.**
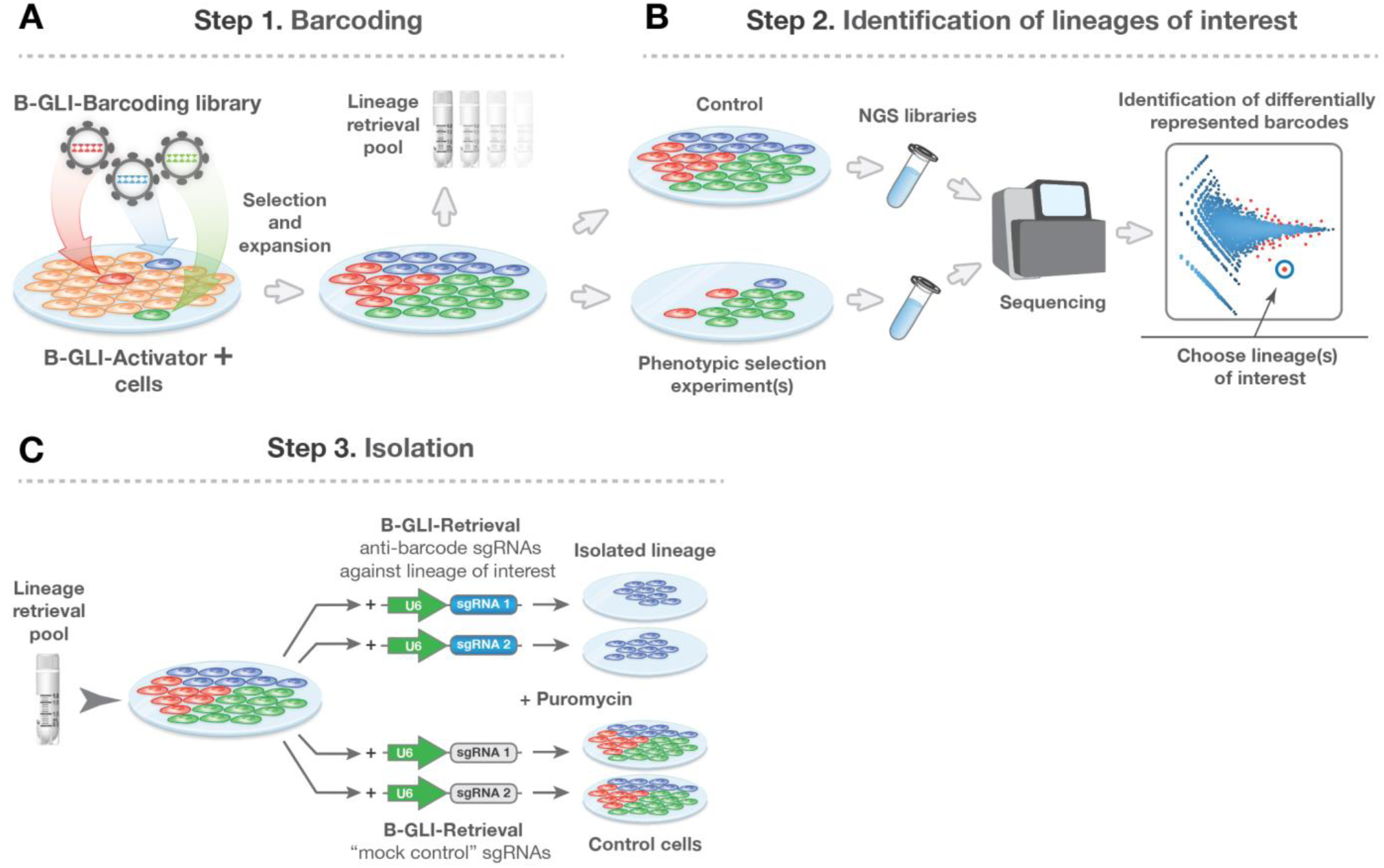
B-GLI schematic workflow. **A**) Cells carrying B-GLI-Activator construct are transduced with lentiviral B-GLI-Barcoding library. The transduced cells are selected with hygromycin, expanded to achieve reasonable lineage representation (e.g. ∼10^2^-10^5^ cells per barcode) and a part of cells is preserved for subsequent lineage isolation (lineage retrieval pool). (**B**) Cells are subjected to a phenotypic selection (e.g. drug treatment, differentiation, tumor initiating frequency), genomic DNA is extracted from the selected cells, and the barcodes are amplified to generate the NGS library. Lineages that exhibit differential phenotype are selected by identification of differentially represented barcodes from count data. (**C**) The preserved fraction of barcoded cells (retrieval pool) is infected with B-GLI-Retrieval lentivirus carrying sgRNAs against the barcodes of interest. In applications where the isolated lineage(s) needs to be compared to the bulk parental cell population, a control cell pool can be produced using B-GLI-Retrieval vector with control sgRNAs.

#### Step 1 – Barcoding

Cell culture is transduced with a virus carrying B-GLI-Activator construct. Then, the lentiviral B-GLI-Barcoding library is delivered to cells, typically at a very low MOI. The transduced culture is expanded and part of the barcoded cell culture is cryopreserved for later use as a cell pool for lineage retrieval (lineage retrieval pool);

#### Step 2 - Identification of the lineages of interest

The barcoded cell culture is assayed to select for cell subpopulation(s) displaying the phenotype of interest. Then NGS libraries are prepared from the genomic DNA of the non-treated and assay-selected cells (e.g., drug-resistant clones, differentiated cells, or metastases) as described in Methods. NGS barcode count data quantification identifies the enriched and depleted barcodes in the assay-selected cells as compared to control pool, pinpointing the differentially-responding subclonal lineages.

#### Step 3 -Isolation of the subclones of interest

Anti-barcode sgRNAs targeting differentially-represented barcodes are delivered to the cryopreserved retrieval cell pool using B-GLI-Retrieval vector. The lineages are isolated by puromycin selection. When needed, control cells are isolated using “mock control” sgRNAs isolation of lineage(s) by puromycin selection and subsequent analyses.

### Controls implemented in the B-GLI system

To control for sgRNA off-target effects (e.g. genome-wide off-targets), we used independent anti-barcode sgRNAs to isolate the same lineage of interest (Figure 1B and 2C). For that, we introduced 2 PAMs into a barcode region which enable lineage isolation with 2 independent anti-barcode sgRNA. Comparison of the independent isolates of the same lineage provides means to confirm that observed lineage-specific properties are not due to sgRNA-specific effects.

To control for possible cellular stress response-related effects of the experimental procedures^35^ (lentiviral infections, expression of the transgenes, and sequential selection with 3 different antibiotics), we equipped the barcoding vector with 2 defined control sgRNA targets common for all the barcoding constructs in the library (Figure 1B and 2C). Recruitment of Cas9-VPR to either of the control sgRNA targets leads to activation of the puromycin resistance gene expression in all the barcoded cells and, therefore, the bulk culture can be “isolated” to provide an equally-treated control for comparing isolated lineages and bulk cell population. Furthermore, additional “mock control” barcodes can be used to verify that all the components of the isolation system function correctly at any point after the barcoding step (Figure 1A).

### Optimization of barcode placement and validation of B-GLI approach

To test the efficacy of the barcode-guided puromycin resistance induction, we first infected Mia-PaCa-2 and OVCAR4 cell lines with B-GLI-Activator vector and the B-GLI-Barcoding vector carrying 2 defined sgRNA targets (Figure 3A and Supplementary Figure 1B). Next, we infected the cells with B-GLI-Retrieval(GFP) vectors that carry either of the targeting sgRNAs or the non-targeting sgRNA, and compared cells sensitivity to puromycin 5 days post-infection. The cells infected with targeting sgRNAs demonstrated high viability upon treatment with increasing doses of puromycin, while the viability of the cells that received non-targeting sgRNA approached zero (Figure 3B). This experiment provided the evidence that anti-barcode sgRNAs assisted the recruitment of dCas9-VPR to the barcodes and enabled lineage-specific activation of puromycin resistance. Also, the results indicate a lack or negligible level of possible leakage of the mCMV promoter upstream of Puro cassette, and therefore, minimal-to-none background puromycin resistance. We did not observe any background resistance due to aberrant activity of mCMV in any of the 5 cell lines that we tested (Figure 3B as a representative experiment and data not shown).

**Figure 3.**
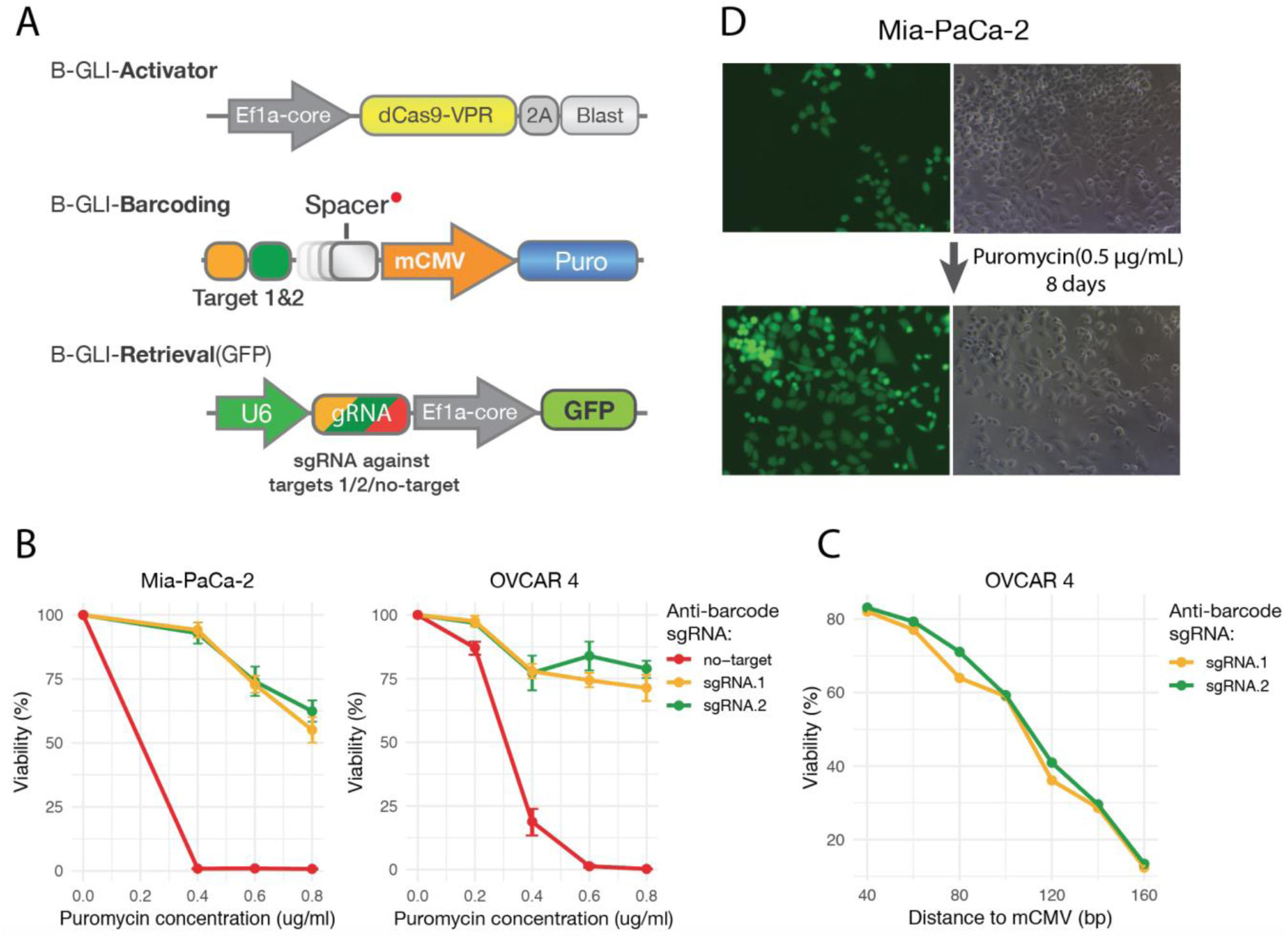
B-GLI system validation and optimization experiments. (**A**) Components of the B-GLI-system used in the validation and optimization experiments. Sequence containing two defined sgRNA targets (Target#1 and Target#2; see Supplementary Figure 1C) were cloned into the barcode region of B-GLI-Barcoding vector. Seven versions of this construct carrying spacer 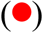 of different length (40, 60, 80, 100, 120, 140, 160 base pairs) were produced as described in Methods to test the optimal barcode position relative to the mCMV promoter. Three B-GLI-Retrieval(GFP) vectors carrying sgRNAs against defined targets (Target#1 and Target#2) and non-targeting sgRNA were produced as described in Methods. (**B**) Mia-PaCa-2 and OVCAR4 (B-GLI-Activator positive) cells were transduced with B-GLI-Barcoding lentivirus carrying defined barcodes and the 40bp spacer. Then, the cells received B-GLI-Retrieval(GFP) virus carrying corresponding anti-barcode sgRNAs. Three days post-transduction the cells were treated with puromycin at the indicated dosage for 5 additional days, after which the cell viability was measured using CellTiter Glo reagent. (**C**) B-GLI-Activator positive OVCAR4 cells were independently transduced with B-GLI-Barcoding lentiviruses harboring the spacer of different lengths downstream of the defined sgRNA targets. Cell viability upon treatment with 0.5 μg/mL Puromycin was measured using CellTiter Glo reagent 5 days after transduction with a B-GLI-Retrieval vector harboring corresponding anti-barcode sgRNA. (**D**) Mia-PaCa-2 (B-GLI-Activator positive) cells were transduced with spacer-optimized B-GLI-Barcoding lentivirus at MOI=0.2 to infect only a fraction of the cells. Then the cells received the B-GLI-Retrieval(GFP) vector and were treated with puromycin (0.5 μg/mL). The phase-contrast and epifluorescence images display the culture at day 0 and day 8 of the puromycin selection.

To achieve an improved robustness in puromycin resistance activation upon the barcode/anti-barcode sgRNA match, we optimized the distance from the barcodes to the mCMV promoter of Puro cassette. We cloned random DNA spacers (Figure 3A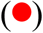) of 40-160 bp between the barcodes cassette and mCMV promoter, and tested the puromycin resistance induction by dCas9-VPR upon the sgRNA delivery. We observed the most potent activation of puromycin resistance when the distance between the barcode region and mCMV was 40 bp (Figure 3C).

To further confirm the optimized system’s capacity to specifically isolate clonal lineages, we delivered B-GLI-Retrieval(GFP) construct carrying sgRNA against defined target (Target#1, Supplementary Figure 1B) only to a fraction of the cells(∼20%). Within days of selection with puromycin, the culture got completely dominated by the cells that received the B-GLI-Retrieval(GFP) construct (Figure 3D), suggesting that the optimized induction of Puro cassette allows specific barcode-guided selection of the cells of interest.

### Full-scale B-GLI experiment

We further challenged the performance of B-GLI system in an experiment aimed at isolating rare lineages from the bulk population. As a model of rare phenotype, we mixed Mia-Paca II cells with and without GFP vector Lego-G2^36^ at 1/50 ratio, respectively. Next, we transduced the mixture with the B-GLI-Activator lentivector and the B-GLI-Barcoding library, and expanded the culture to achieve the barcode representation of ∼100 cells per barcode. To identify the lineages that belonged to GFP+ population, the gDNA from the flow cytometry-sorted GFP-cells and from the non-sorted cell mixture was analyzed by NGS as described in Methods.

To isolate the GFP+ cells from the original barcoded mixture using B-GLI, two depleted barcodes of reasonably large-sized (∼0.1%) lineages were selected to generate B-GLI-Retrieval constructs with anti-barcode sgRNAs. The NGS analysis of the gDNA from the isolated populations confirmed the lineages purity (85% and 98% for lineages 1 and 2, respectively) and, as expected, the isolates were GFP positive (Figure 4B).

**Figure 4.**
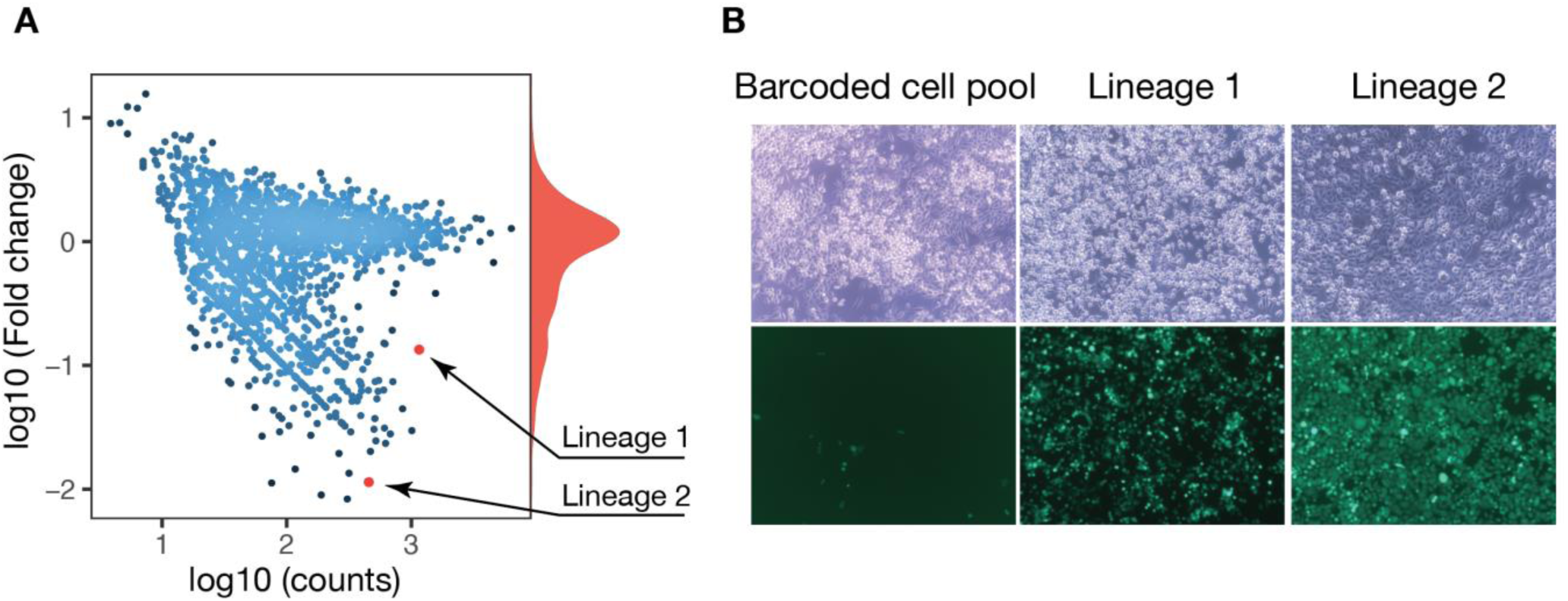
Identification and isolation of phenotypically distinct subclonal lineages in a model experiment. **(A)** The Mia-PaCa-2 culture containing 2% of GFP expressing cells was transduced with B-GLI-Activator and B-GLI-Barcoding library and expanded to achieve the barcode representation of ∼100 cells per barcode. The genomic DNA both from the FACS-sorted GFP-negative cells and from the non-sorted cell mixture was extracted and the sequencing library for NGS was prepared as described in Methods. MA-plot displays differentially represented barcodes in GFP negative Mia-PaCa-2 cells versus bulk cell pool as quantified from the NGS data. Barcode count data was median-normalized using DESeq2 built-in function (estimateSizeFactorsForMatrix). Lineages selected for isolation are marked in red. (**B)** Phase contrast and epifluorescence images of barcoded retrieval pool of Mia-PaCa-2 cells and the isolated lineages.

## Discussion

We have developed B-GLI system as a novel method for specific and controlled isolation of subclonal lineages from heterogeneous cancer cell cultures. The validation experiments indicated that the performance of B-GLI makes it a prospectively useful approach to address the biology of individual lineages via their isolation, expansion, and phenotypic profiling. To our knowledge, methods for DNA barcode-based lineage isolation have been limited so far to CRISPRa-based COLBERT^37^ and Cas9-mediated barcode editing-based SmartCodes^38^. While COLBERT, SmartCodes, and B-GLI share a similar concept of functionalized DNA barcodes for isolation of individual subclones from a heterogeneous culture, B-GLI approach implements a number of improvements (Table 1) that make it applicable to a broader range of research questions.

**Table 1.** Comparison of the existing methods for the barcode-guided clonal lineage isolation.

|  | **COLBERT**<sup>37</sup> | **SmartCodes**<sup>38</sup> | **B-GLI** |
| --- | --- | --- | --- |
| Controls for sgRNA-specific effects | No | Yes (only via genomic off-target prediction algorithms; no direct control) | Yes (direct control via independent isolations with multiple sgRNAs) |
| Control for isolation-associated treatment effects | No | No | Yes (mock isolation via control sgRNAs) |
| Library complexity potential (number of unique barcodes) | High($\sim 10^6$-$10^8$) | Low($\sim 5 \times 10^4$) | High ($\sim 10^6$-$10^8$) |
| Delivery of the isolation system | Cationic lipid transfection | Retroviral transduction | Lentiviral transduction |
| Lineage selection marker | GFP/other | Puromycin/other | Puromycin/other |
| Factors affecting lineage retrieval efficacy | sgRNA on-target efficacy, transfection efficacy | sgRNA on-target efficacy, barcode mutation outcome, transduction efficacy | sgRNA on-target efficacy, transduction efficacy |

The specificity of the isolation method affects the purity of the isolated lineages and hence the accuracy of their downstream profiling. The cross-lineage contamination could arise due to anti-barcode sgRNA targeting barcodes other than the barcode of interest. Promiscuous targeting of multiple barcodes by the same sgRNA is expected to be a relatively rare event due to the high specificity of CRISPR systems; nevertheless, B-GLI offers an opportunity to control for promiscuous targeting by using 2 or more redundant lineage-specific barcode targets for isolation. Yet another possible cause for low specificity is “leakage” of the selection marker expression. Using the B-GLI system we were unable to detect any leakage in any of the 5 cell lines we tested (Mia-PaCa-2, HEK293, KURAMOCHI, OVCAR4, OVCAR5), but if the issue occurs, it should be possible to significantly reduce or eliminate the leakage with longer puromycin selection.

To control for possible biologic effects introduced during isolation-associated procedures throughout the experiment (e.g. infections and antibiotics treatment), B-GLI relies on mock isolation via control sgRNA targets (control barcodes, Figures 1 and 2). To control for potential sgRNA-specific effects (e.g. genomic off-targets), we suggest to compare the lineages isolated via multiple sgRNA targets for each lineage. SmartCodes and COLBERT provide no means for direct control for sgRNA-specific effects and isolation-associated procedures, although the SmartCodes address sgRNA off-target issue via the design of sgRNAs with low predicted probability of off-target. However, the design of SmartCodes systems could be improved by including controls similar to those used in B-GLI system.

The existing lineage isolation methods differ in their factors affecting the lineage retrieval efficiency (the fraction of cells of the lineage of interest that gets isolated after sgRNA delivery and selection, see Table 1). The means of vector delivery commonly limit the efficiency of the lineage retrieval in all three methods. The B-GLI system is fully lentiviral, that is arguably advantageous compared to retroviral infection (used in Smartcodes) and cationic lipid transfection (used in COLBERT for the 2 out of 3 vectors). Transfection could be useful in certain hard-to-transduce cell types, but lipofection of large cell pools is costly, and can be detrimental to cells. Additionally, efficiency of the lineage isolation by COLBERT^37^ is limited by the throughput of flow cytometry-based sorting, especially when isolating rare subpopulations. Nevertheless, our current experimental data do not enable to conclude which of the systems provides the best lineage retrieval efficiency, and this performance measure remains to be compared.

Starting the lineage tracing experiment with barcoding of an adequate number of cells is important to avoid a sampling bottleneck when the experimental question addresses subpopulation heterogeneity. High complexity of the DNA barcode library allows one to tag a large number of cells avoiding non-uniquely labeled clones. THe complexity potential of B-GLI and COLBERT DNA barcoding libraries is comparable to the existing libraries used for lineage tracing (∼10^6^-10^8^ unique barcodes)^13,39^, and they are limited only by the number of random bases in the barcode and by the cloning/transformation efficacy. The complexity of SmartCodes system, however, is limited by the strict sgRNA design rules and scalability of pooled oligo synthesis^40^ and therefore it achieves the complexity of 5*10^4^ individual barcodes^38^. Such a moderate complexity will affect the number of uniquely labeled cells. On estimation, barcoding of 10 000 cells using a library of such complexity will result in about 20% of non-uniquely labeled cells^27^, which leads to merging of signal from the lineages of different origin carrying the same barcode. The B-GLI and COLBERT systems are easily scalable allowing for high complexity DNA barcoding to enable tracing of millions of individual lineages. For the experiments presented in the current manuscript, we used a library of 5*10^6^ unique DNA barcodes, which would result in ∼0.2% of non-uniquely labeled cells if 10000 cells are barcoded. We conclude that B-GLI barcoding library can be used for high-throughput lineage tracing experiments, regardless of the isolation procedure, thanks to its complexity.

Profiling of the differences between specific cancer cell lineages allows dissecting and exploring the biological basis of cellular plasticity and clonal evolution in cancer. The isolation of individual lineages extends the use of different molecular profiling methods beyond genomics to monitor and follow-up the regulatory changes (e.g. epigenetic modifications and kinome reprogramming) accompanying the clonal expansion in unperturbed and perturbed conditions. Furthermore, for genomic analyses, isolation and expansion of the subclonal lineages of interest will additionally lower the costs for sequencing, since less deep NGS is needed to define mutation-driven phenotypes in pre-isolated enriched subclonal populations. Indeed, such a setup is not limited to cancer biology. Broadly speaking, coupling DNA barcoding with a robust lineage isolation method like B-GLI, is a gateway for the studies of dynamic changes in the heterogeneous, evolving cellular systems, such as cancer, regenerating tissues, differentiating cells, or other similar imaginable models with a higher level of resolution.

## Methods

### Generation of B-GLI vectors

#### B-GLI-Activator

Lenti-dCas9-VP64-Blast^41^ (Addgene plasmid #61425) was amplified with primers C5-dCas9VP64.F and C5-dCas9VP64.R (see table) to cut out the VP64 activator and attach AarI cloning overhangs compatible with the Golden Gate cloning method^42^. The VPR activator was amplified from SP-dCas9-VPR^32^ (Addgene plasmid #63798) with C5-sp.dCas9VPR.F and C5-sp.dCas9VPR.R. PCR fragments were purified and ligated using the Golden Gate (AarI) cloning approach (see Supplementary methods for details).

#### B-GLI-Barcoding

To generate B-GLI-Barcoding construct, we cloned the barcode cloning site, mCMV promoter and puromycin resistance gene into a previously generated acceptor vector LG-EF1a-Hygro (LentiGuide-Puro vector^43^ (Addgene plasmid #52963) where U6-sgRNA cassette was removed and the Puro cassette replaced with Hygro). The barcode cloning site and the mCMV promoter were ordered as a single double-stranded DNA from Integrated DNA Technologies (IDT) as a gBlocks® gene fragment (see Table for sequence). The puromycin resistance gene was amplified from LentiGuide-Puro^43^ (Addgene plasmid #52963) with Puro.frLG-forVec15.3.F and Puro.frLG-forVec15.3.R primers (Supplementary Table 1). The acceptor vector (LG-EF1a-Hygro) was amplified with LG-MU6-pos16-F and LG-MU6-for.CS.mCMV-R primers (Supplementary Table 1). The three fragments were circularized using the Golden Gate (Esp3I) protocol^42^ (see Supplementary methods).

#### B-GLI-Retrieval

We generated the B-GLI-Retrieval vector by replacing the sgRNA scaffold in LentiGuide-Puro^43^ to an optimized sgRNA^E-F^ scaffold^34^ to obtain LentiGuide-Puro-EF vector. LentiGuide-Puro-EF was amplified to get a linear backbone for cloning of Hygro (for B-GLI-Retrieval-Hygro) or eGFP (for B-GLI-Retrieval-GFP) under the control of an Ef1a-intron promoter. For that, eGFP was amplified from the LEGO-G2^36^ vector (Addgene plasmid #25917) with eGFP-LEGO-in2A1E.ef1.F and eGFP-LEGO-in2A1E.ef1.R primers (Supplementary Table 1). A hygromycin resistance gene was amplified from pLenti PGK Hygro DEST^44^ (addgene plasmid #19066) with Hygro-into-LG-MU6C-PuroOut.F and Hygro-into-LG-MU6C-PuroOut.R primers (Supplementary Table 1). Fragments were ligated using the Golden Gate protocol (AarI; see Supplementary methods).

All PCR amplifications were performed with high-fidelity SuperFI polymerase (Thermo, cat. 12351010). For the long templates (>5 kb), the elongation time was set to 1.5 min/kb. We used 1 M betaine (SigmaAldrich, cat. 61962-50G) as a PCR enhancer. We used a high starting amount (50 ng) of the template to limit the number of amplification for each copy of the template, thereby reducing mutagenesis. Golden Gate reactions were transformed into NEB® Stable Component *E. coli* (NEB, cat. C3040H) according to the manufacturer’s instructions. The identity and correctness of all resulting vectors were verified by Sanger sequencing and functional tests.

### Barcodes library cloning into B-GLI-Barcoding vector

Barcode template (Barc.LGMU6.templ; see Supplementary Table 2) was ordered as a single-stranded DNA from SigmaAldrich. The barcode was amplified with Barc.LGMU6.aarI.ampl.F and Barc.LGMU6.aarI.ampl.R (Supplementary Table 1) to include AarI-compatible cloning overhangs. Five microliters of the reaction was transferred to a new 50 µl PCR reaction with an excess of Barc.LGMU6.aarI.ampl.F and Barc.LGMU6.aarI.ampl.R primers (Supplementary Table 1). The reaction was run for one cycle to produce double stranded barcodes with no mismatches. The barcodes were purified using AMPure XP SPRI beads (Beckman Coulter, cat. A63880) according to the manufactirer’s instructions. The barcodes were cloned into B-GLI-Barcoding vector using the Golden gate method (AarI; see Supplementary methods). To reduce possible contamination with original circular B-GLI-Barcoding vector, extra 2 µl of AarI was added to the reaction mix at the end of the Golden Gate reactionfollowed by overnight incubation at 37°C.. The reactions were magnetic bead-purified (Beckman Coulter, cat. A63880) and treated with Plasmid-Safe(tm) ATP-Dependent DNase (Lucigen, cat. E3101K). The reaction was again concentrated by magnetic bead purification and transformed into Lucigen Endura(tm) electrocompetent *E. coli* (Lucigen, cat. 60242-2) using Bio-Rad MicroPulser Electroporator (cat. #1652100) according to the manufactirer’s instructions. The transformation reactions were plated onto five 15 cm LB-agar dishes with 100 µg/ml ampicillin and incubated for 16 h at 32°C. Bacteria were collected and the plasmid DNA was purified using NucleoBond® Xtra Midi kit (MACHEREY-NAGEL, cat. 740410.50). The transformation efficiency was estimated by plating 1/10000 of the transformation reaction onto 15 cm LB-agar plate with 100 µg/ml ampicillin and counting number of clones after incubation overnight at 37°C.

### Cloning sgRNA into B-GLI-Retrieval vector

sgRNA design and cloning was performed as described^45^ except for the Golden Gate cloning step (see protocols for Esp3I Golden Gate protocol in the Supplementary methods).

### Cell culture

Mia-PaCa-2 cell line was maintained in DMEM (Dulbecco’s Modified Eagle Medium; Gibco(tm), catalog number: 41965039), and OVCAR4, OVCAR5, and KURAMOCHI ovarian cancer cell lines were cultured in RPMI 1640 (Gibco, cat. 11875093) in a humidified incubator at 37°C with 5% CO2. The media were supplemented with 10% Fetal Bovine Serum (FBS, Gibco, cat. 26140079), Penicillin-Streptomycin (100 U/mL, Gibco, cat. number: 15140122), 2 mM Glutamine (Thermo, cat. number 25030081). HEK293FT cells were cultured in DMEM with 10% FBS, 100 U/mL Penicillin-Streptomycin, 2 mM Glutamine and 20 mM HEPES. All the cell lines were from ATCC.

### Lentiviral packaging and transduction

HEK 293FT cells were seeded in a density of 10^5^ cells/cm^2^. Next day the cells were transfected with a transfer plasmid, packaging plasmids pCMV-VSV-G^46^ (Addgene plasmid #8454) and pCMV-dR8.2 dvpr^46^ (Addgene plasmid #8455) using Lipofectamine 2000 (Thermo, cat. 11668019) transfection reagent according to the manufacturer’s instructions. Virus supernatants were collected 48 h post-transfection. The titer of the virus was determined as described^47^.

For transduction, the cells were plated at 5*10^4^ cells/cm^2^ and the volume of viral supernatant containing the desired number of lentiviral particles was added to culture medium in presence of 8 μg/mL polybrene (Sigma, cat. 28728-55-4). The medium was changed to the standard culture medium 24 h post-transduction.

### Validation and optimization experiments

A DNA sequence with two defined sgRNA targets (Target#1 and Target#2) were selected to have a reasonably high on-target activity according to the sgRNA design rules^33^ (efficiency score of around 0.8 as predicted by Benchling[Biology Software]. (2019). The B-GLI-Barcoding vector with sgRNA targets and 40 bp spacer was generated by cloning a single insert into a barcode cloning site using Golden Gate cloning (AarI; see Protocols in the Supplementary methods). The insert was produced by overlap extension PCR using Vec15.Ins.VPR.11 and Vec15.Ins.VPR.12 oligos (Supplementary Table 1). Vectors with spacer lengths of 60-160 bp were generated by simultaneous cloning of two inserts. The first insert is common for all the constructs. It was generated by an overlap extension PCR using Vec15.Ins.VPR.11 and Vec15.Ins.VPR.13 (Supplementary Table 1). The second insert determines the length of the spacer and, therefore, specific for each construct. The inserts were generated by PCR-amplification of RRE-oligo (Supplementary Table 2) forward primers Vec15.Ins.VPR.X1, where X is a number indicating the spacer length (2 - 60 bp; 3 - 80 bp; 4 - 100 bp; 5 - 120 bp; 6 - 140 bp; 7 - 160 bp) and reverse primer Vec15.Ins.VPR.22 (Supplementary Table 1). The inserts were cloned using the Golden Gate protocol (AarI, see Protocols in the Supplementary methods). Corresponding sgRNAs were cloned into a B- GLI-Retrieval(GFP) vector using oligos Vec15.test.g1-F, Vec15.test.g1-R for sgRNA Target #1 and Vec15.test.g2-F, Vec15.test.g2-R for sgRNA Target #2 (Supplementary Table 1).

### Full-scale B-GLI experiment

Mia-PaCa-2 GFP positive cells were made by transduction Mia-PaCa-2 cells with virus particles carrying LEGO-G2 vector at MOI of ∼10. Mix of Mia-PaCa-2 and Mia-PaCa-2 GFP+ cells were transduced with lentiviral particles generated using B-GLI-Barcoding library with a complexity of ∼5*10^6^ barcodes. After incubating 24 h with virus, cells were selected with 150 µg/ml hygromycin for 7 days. We used a MOI of <0.01 to infect 10^6^ cells to get ∼10000 barcoded cells. Using low MOI ensures that most of the cells receive only one barcode. After selection, cell lines were expanded to achieve a representation of ∼100 cells per barcode. To improve the viability of the culture during selection and expansion, the cells were kept at a density higher than 3*10^4^ cells/cm^2^. After expansion, a part of the barcoded cells were frozen to generate a retrieval pool. The rest of the cells were sorted for a GFP negative subpopulation using a Sony SH800Z Cell Sorter. Generation of NGS libraries and count data analysis were performed as described below.

Two lineages depleted in the GFP-negative pool were selected for isolation. One anti-barcode sgRNA target for each lineage was selected. Oligos for sgRNA cloning were ordered from SigmaAldrich. The sgRNAs were cloned into the B-GLI-Retrieval(Hygro) vector as described (see methods). Virus titers were evaluated in the wild-type Mia-PaCa-2 cells. The retrieval pool was thawed and infected with the B-GLI-Retrieval vectors at a MOI of ∼2. Four days after infection, lineages were isolated by 0.5 µg/ml puromycin treatment for 8 days. NGS library was prepared from isolated lineages and sequenced to confirm lineage purity.

### NGS library preparation and sequencing

Genomic DNA was isolated using NucleoSpin® Tissue kit (MACHEREY-NAGEL) following manufacturer instructions. Barcode cassette was amplified from genomic DNA with primers P5.seq-B-GLI.v1 and P7.seq-B-GLI.v1 using OneTaq® DNA Polymerase (NEB, cat. M0480; see Protocols in the Supplementary methods). PCR reactions were purified with NucleoSpin® Gel and PCR Clean-up (MACHEREY-NAGEL). Purified reactions were PCR-amplified with primers Illumina_indX_F and Illumina_indX_R (where XXXXXXXX is index) to add sample-specific indexes (for sample multiplexing) and Illumina adapters. The second round PCRs were performed with NEBNext® Ultra(tm) II Q5® Master Mix (NEB, cat. M0544). Amplified samples were purified with AMPure XP SPRI beads (Beckman Coulter, cat. A63880). NGS library were sequenced on HiSeq2500 sequencer (Illumina) with 100 bp paired-end sequencing (10% PhiX DNA spike-in was used). We added a 15 bp random sequence in the P5.seq-B-GLI.v1 primer to increase sequence diversity which is intended to improve cluster calling.

### Barcode calling from NGS data

To reconstruct original barcodes by accounting for PCR and sequencing errors, we developed a custom Python script (to be posted at GitHub soon). Raw sequences from FASTQ files were first processed to extract sequences in between the barcode flanking sites (ACCGCGGC and TTCACCCG). The extracted sequences with length not equal to 41 bp were discarded. Then, the barcode sequences were filtered according to the minimal (>20) and average (>30) Phred quality scores, and the arcode frequencies were calculated. To account for PCR and NGS errors we adopted an algorithm similar to previously reported^13,48^. Briefly, the algorithm starts by picking a barcode with the lowest frequency and calculates Hamming distances to all the other barcodes. If there is a similar barcode with the Hamming distance lower than the defined threshold (threshold depends on the barcode length, for 41 bp barcode length we empirically derived a threshold of 4 bases) then the counts of the picked barcode is added to the found similar barcode. When there are several similar barcodes having the same Hamming distance, the counts of a picked barcode are added to the barcode with the highest frequency (unlike algorithm applied in Thielecke et. al.^13,48^). The algorithm is iteratively applied to the sorted barcode list starting from the the barcode with the lowest count.

### Barcode count data analysis

Barcode count data was analyzed using a custom R script. Read counts were median-normalized by estimating size factors for each sample using built-in DESeq2 function “estimateSizeFactorsForMatrix”. Then each column was divided by the estimated size factor. To identify depleted barcodes we used log2 read count fold changes between unsorted and sorted cell populations.

