## Supplementary File for "DNA barcode-guided lentiviral CRISPRa tool to trace and isolate individual clonal lineages in heterogeneous cancer cell populations"

### Supplementary methods

#### Golden gate protocol (for AarI OR Esp3I):

| | |
|---|---|
| Rapid Ligation Buffer (Thermo, cat. K1422) | 4 µl |
| AarI (2U/ul) (Thermo, cat. ER1581) OR<br>Esp3I FastDigest (Thermo, cat. FD0454) | 1 µl |
| T4 DNA Ligase (5 U/µL) (Thermo, cat. EL0014) | 0.25 µl |
| linear fragments to be ligated (with restriction enzyme sites overhangs) | 50-500 ng |
| Oligo (only for use with AarI; provided with the AarI enzyme) | 0.4 µl |
| ddH2O | up to 20 µl |

#### Golden gate cycling conditions

| Temperature | Time | Cycles |
|---|---|---|
| 37C | 5 min | 1 |
| 37C<br>22C | 5 min<br>3 min | 50 |
| 37C | 10 min | 1 |
| 65C | 15 min | 1 |
| 4C | hold | 1 |

#### Barcode PCR-amplification protocol

| Round 1 | | Round 2 | |
|---|---|---|---|
| **Component** | **Amount (µl)** | **Component** | **Amount (µl)** |
| Standard Buffer (5x) | 10 | NEBNext® Ultra™ II Q5® Master Mix | 10 |
| dNTP | 1 | Illumina_indX_F(10µM) | 1 |
| P5.seq-B-GLI.v1(100µM) | 0,25 | Illumina_indX_R(10µM) | 1 |

| | | | |
|---|---|---|---|
| P7.seq-B-GLI.v1(100µM) | 0,25 | Amplified barcodes (from Round 1; purified;10-100 ng) | 2 |
| Betaine (5M) | 10 | $H_2O$ | 6 |
| genomic DNA (up to 4µg) | 20 | | |
| OneTaq polymerase | 0,25 | | |
| $H_2O$ | 8,25 | | |

| Cycling conditions | | | | | |
|---|---|---|---|---|---|
| **Round 1** | | | **Round 2** | | |
| Temperature | Time | Cycles | Temperature | Time | Cycles |
| 94 ºC | 10 min | 1 | 98 ºC | 1 min | 1 |
| 94 ºC | 30 sec | 24 | 98 ºC | 20 sec | 4-8 |
| 55 ºC | 60 sec | | 72 ºC | 40 sec | |
| 68 ºC | 80 sec | | 68 ºC | 80 sec | |
| 68 ºC | 2 min | 1 | 68 ºC | 2 min | 1 |

#### Supplementary Table 1. Primers

| | |
|---|---|
| C5-dCas9VP64.F | TAATTACACCTGCTCGTGTTTATTAACTGTACAGGCAGTGGAGAGG |
| C5-dCas9VP64.R | ATAATACACCTGCTATATAATCAGCATGTCCAGGTCGAAATCATCA |
| C5-sp.dCas9VPR.F | TAATTACACCTGCTGCAATTAACTCTAGAAGTTCCGGATCTCCG |
| C5-sp.dCas9VPR.R | AATATACACCTGCCTCAAAACAGAGATGTGTCGAAGATGGACAG |
| Puro.frLG-forVec15.3.F | GTACTGCGTCTCTCACCATGACCGAGTACAAGCCCACG |
| Puro.frLG-forVec15.3.R | CGACAGCGTCTCATCAGGCACCGGGCTTGC |
| LG-MU6-pos16-F | GTACTGCGTCTCTCTGAGATAAAGTTTTAAACAGAGAGGAATCTTTG |
| LG-MU6-for.CS.mCMV-R | CAGCGTCTCAGTGTCATCTTTGCACCCGGG |
| eGFP-LEGO-in2A1E.ef1.F | TAATTACACCTGCTGCAATGAAAAAGCCTGAACTCACCGCG |

| | |
|---|---|
| Hygro-into-LG-MU6C-PuroOut.R | AATATACACCTGCCTCAGCGTCTATTCCTTTGCCCTCGGACGAG |
| Hygro-into-LG-MU6C-PuroOut.F | TAATTACACCTGCTGCAATGAAAAAGCCTGAACTCACCGCG |
| Barc.LGMU6.aarI.ampl.F | TAGCTGCACCTGCTGCAATCGTCTCCTCGGAGAGATAACCGC |
| Barc.LGMU6.aarI.ampl.R | TGTTAACACCTGCAATGACCTTCAGGTCCGAGTCGGGTGAA |
| Vec15.Ins.VPR.11 | TCCTTACACCTGCTCGTATCGGGGTGCCATAACGAATACACGGTGG CCACCTCGGAGAGATAACCGG |
| Vec15.Ins.VPR.12 | ATGCTACACCTGCTATAACCTCCCCGGTTATCTCTCCGAGGTGG |
| Vec15.Ins.VPR.13 | ATGCTACACCTGCTATAGTGGCCCCGGTTATCTCTCCGAGGTGG |
| Vec15.Ins.VPR.21 | CTATCGCACCTGCTCGTCCACGGAGACCTAGGACCATTCCA |
| Vec15.Ins.VPR.22 | AGGTAACACCTGCTATAACCTTGGAATGGTCCTAGGTCTCC |
| Vec15.Ins.VPR.31 | CTATCGCACCTGCTCGTCCACTACCAGGCCTACGGTGG |
| Vec15.Ins.VPR.41 | CTATCGCACCTGCTCGTCCACGTTTCCTCTTGGATACCTGGTACC |
| Vec15.Ins.VPR.51 | CTATCGCACCTGCTCGTCCACGACCGAAGCGCGGC |
| Vec15.Ins.VPR.61 | CTATCGCACCTGCTCGTCCACGGAACACCCACACTCCG |
| Vec15.Ins.VPR.71 | CTATCGCACCTGCTCGTCCACAGCATCGGGCCCGTA |

#### Supplementary Table 2. dsDNA/oligos

| | |
|---|---|
| Barc.LGMU6.templ (ssDNA) | TCTCCTCGGAGAGATAACCGCGGCNNNNNNNNNNNNNNHNHNVVRN GGHNNNNNNNHNHNVVRNGGHTTCACCCGACTCGGACCTGA |
| gBlock-CS(AarI)-mCMV-esp3I (dsDNA) | GTACTGCGTCTCTACACCCACACTCCGGTGGACCGAAGCGCGGCAC CAGGGTTTCCTCTTGATCGGCATGCAGGTGTACGTATTGCTATGTGC GCACCTGCTTCGAGGTGCACCGGCGCCGGCCATCGGCCTTGGAGGC CGTGGTAGGATAGGCGTGTACGGTGGGAGGCCTATATAAGCAGAGC TCGTTTAGTGAACCGTCAGATCGCCTGGAGACGCCATCCACGCTGTT TTGACCTCCATAGAAGACACCGGGACCGATCCAGCCTCTCGACATT CGTTGGATCCGCCACCTGAGACGCTGTCG |
| RRE-oligo | AGCGGCGGCTAACCAAGGCGCCTGCCACGGCAGCAGCATCGGGCC CGTACAAAGGGAACACCCACACTCCGGTGGACCGAAGCGCGGCAC |

| | |
|---|---|
| | CAGGGTTTCCTCTTGGATACCTGGTACCAGGCCTACGGTGGCCTGGAGACCTAGGACCATTCCA |
| Vec15.test.g1-F | CACCGTGCCATAACGAATACACGG |
| Vec15.test.g1-R | AAACCCGTGTATTCGTTATGGCAC |
| Vec15.test.g2-F | CACCGCCACCTCGGAGAGATAACCG |
| Vec15.test.g2-R | AAACCGGTTATCTCTCCGAGGTGGC |

**Supplementary Table 3. Primers for NGS library preparation**

| | |
|---|---|
| P5.seq-B-GLI.v1 | CCCTACACGACGCTCTTCCGATCTNNNNNNNNNNNNNNNNCTTGATCGTCTCCTCGGAGAG |
| P7.seq-B-GLI.v1 | GTGACTGGAGTTCAGACGTGTGCTCTTCCGATCTCCTTCAGGTCCGAGTCGGGTGAA |
| Illumina_indX_F | AATGATACGGCGACCACCGAGATCTACACXXXXXXXXACACTCTTTCCCTACACGACGCTCTTCCGATC*T |
| Illumina_indX_R | CAAGCAGAAGACGGCATACGAGATXXXXXXXXGTGACTGGAGTTCAGACGTGTGCTCTTCCGATC*T |

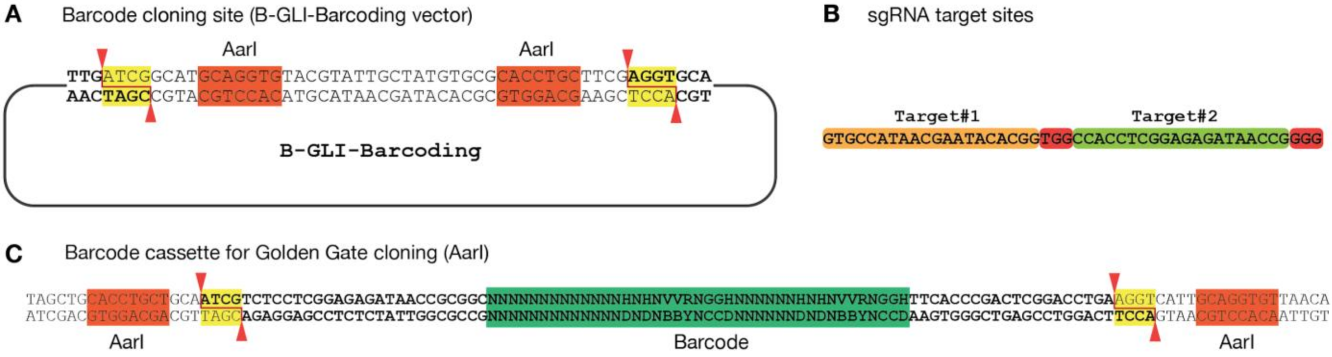


**Supplementary figure 1. (A)** Barcode cloning site of B-GLI-Barcoding vector. AarI recognition sites are marked in red; AarI cut sites are marked in yellow. **(B)** sgRNA target sites as used in validation and optimization experiments. PAM sites are marked in red **(C)** Amplified barcode cassette as used for barcode cloning into B-GLI-Barcoding vector.
